# Understanding the LLM-based gene embeddings

**DOI:** 10.64898/2025.12.19.695582

**Authors:** Yufei Cai, Dailin Gan, Hui Zhang, Jun Li

## Abstract

Large language model (LLM)–derived gene embeddings, generated from brief NCBI gene descriptions, have shown strong performance in recent biological applications, yet the biological information they contain remains unclear. We evaluate these embeddings using a Gene Set Enrichment Analysis (GSEA)–based framework that treats each embedding dimension as a potential carrier of pathway-level information. OpenAI’s embeddings recover over 93% of Hallmark and C2 pathways, with pathway signals distributed across many dimensions. Even embeddings generated from gene symbols alone recover more than 64% of pathways, indicating substantial prior biological knowledge embedded in the model. Comparing 11 small language models reveals that domain-specific models perform best with minimal input, but all models approach OpenAI-level coverage when given modest textual context. Collectively, these results show that LLM-derived embeddings encode unexpectedly extensive pathway-level information, supporting their use as lightweight, informative representations for downstream biological analysis.

## Introduction

Transformer-based large language models (LLMs), originally developed for natural language processing, are now being repurposed to analyze biological data, offering new avenues for understanding complex biological systems. In particular, their integration into gene-expression data analysis has garnered substantial interest due to their remarkable predictive capabilities across diverse applications in fundamental biological problems (see, e.g., [1–7] for a review).

Current methods for integrating LLMs into gene-expression analysis can be broadly categorized into three main approaches. The first approach leverages the question-answering capabilities of chatbot models like ChatGPT. For instance, Hou and Ji [8] utilized ChatGPT to identify cell types by providing the top ten differentially expressed genes for each cell type and directly asking the chatbot to classify the cell. While this strategy is straightforward and effective for certain applications, it suffers from key limitations. Specifically, ChatGPT’s difficulty in processing and understanding numerical data, coupled with the challenge of designing effective prompts for complex tasks, restricts its applicability to more complex problems, particularly those involving numerical data.

The second approach involves training transformer-based foundation models specifically for gene-expression data. Examples include scBERT [9], Geneformer [10], scGPT [11], and scFoundation [12]. These models are trained on extensive collections of single-cell RNA-seq data, typically encompassing thousands of datasets and millions of cells. Once trained, they can be fine-tuned on smaller, user-specific datasets to suit particular applications. While this method offers substantial predictive power in certain applications, its resource-intensive pre-training is inaccessible to most research groups [13], and its performance after fine-tuning often depends sensitively on how well the biological conditions represented in the pre-training data align with those of the target application—an issue that has contributed to the mixed performance reported in recent evaluations [14].

The third approach, pioneered by GenePT [15], circumvents the need for training foundation models from scratch by utilizing general-purpose LLMs such as ChatGPT. Specifically, GenePT employs OpenAI’s text-embedding function [16] to transform standard NCBI gene descriptions [17] into long, dense numeric vectors known as embeddings. These embeddings are presumed to encapsulate the semantic content of the gene descriptions (e.g., functional roles, associated processes, or molecular characteristics), and they can be directly used in downstream tasks. For example, the GenePT method demonstrated effective cell type labeling using these embeddings, and a subsequent method, scHOTT [18], further optimized this process. More importantly, these embeddings have been successfully applied to solve more complex biological problems, such as predicting the outcomes of gene perturbation experiments, yielding substantial gains in modeling efficiency [19–21]. Related work has further incorporated GenePT-style representations—either directly or as a source of in-spiration—into broader multimodal frameworks that integrate gene-expression data with spatial transcriptomics or histopathological image features, supporting applications such as spatial domain identification, cell-state characterization, and disease-related prediction [22–25]. Notably, the same embedding paradigm has also been generalized beyond genes to other biomedical entities, such as clinical concepts in electronic health records, where LLM-derived embeddings of concept descriptions have been used for disease risk prediction [26, 27].

Among these three approaches, the embedding-based strategy offers a particularly practical and versatile complement to the others. The text-embedding service of ChatGPT is highly cost-efficient—substantially cheaper than the question-answering API—and recent studies have shown that it can often be replaced by free, open-source LLMs without a notable loss in performance [28]. Moreover, embeddings provide a flexible numerical representation of genes, making them compatible with a wide spectrum of machine-learning techniques, from traditional methods such as logistic regression and random forests to modern deep-learning architectures. These features make the embedding-based approach broadly accessible and easy to integrate into diverse analytical pipelines, while remaining complementary to the strengths of the other two approaches, which excel in different problem settings.

However, the advantages of this third approach rest on a critical assumption: that the embeddings, as numeric vectors, encapsulate rich information about the genes they represent. While the success of various applications using these embeddings suggests the validity of this assumption, it has never been directly validated. In this work, we rigorously investigate this assumption by exploring the extent to which these embeddings capture biologically meaningful information about genes. In particular, we assess this by examining whether genes belonging to the same biological pathways exhibit coherent structure within the embedding space.

## Methods

### NCBI Text Descriptions Do Not Automatically Imply Information-Rich Embeddings

This subsection illustrates why short NCBI descriptions cannot be assumed to yield information-rich embeddings and therefore motivates the analyses presented in the subsequent sections.

Figure 1 (A) and (B) present the NCBI descriptions of two well-studied human genes, *TP53* and *BRCA1*. *TP53*, often referred to as the “guardian of the genome,” encodes a tumor suppressor protein that plays a critical role in regulating the cell cycle, DNA repair, and apoptosis. Mutations in *TP53* are associated with a broad spectrum of cancers. Similarly, *BRCA1* encodes another tumor suppressor protein, primarily involved in DNA repair. Mutations in *BRCA1* are strongly linked to an elevated risk of breast, ovarian, and other cancers, making it a key target for genetic testing and targeted therapies. Both *TP53* and *BRCA1* are foundational to cancer research and clinical care, having been the focus of extensive investigations that have revealed a wealth of knowledge about their regulation, interactions with other genes, and roles in cancer biology. However, the NCBI descriptions of *TP53* and *BRCA1* are relatively brief. Each gene is described in a single paragraph that, while touching upon multiple aspects of the gene’s function, lacks depth and detail. These descriptions provide a high-level overview but do not comprehensively capture the genes’ regulatory networks, interactions, or pathway involvement—key aspects crucial for understanding their biological roles. Importantly, they are not curated to represent regulatory or pathway-level knowledge, making them an imperfect source for downstream inference.

**Figure 1:**
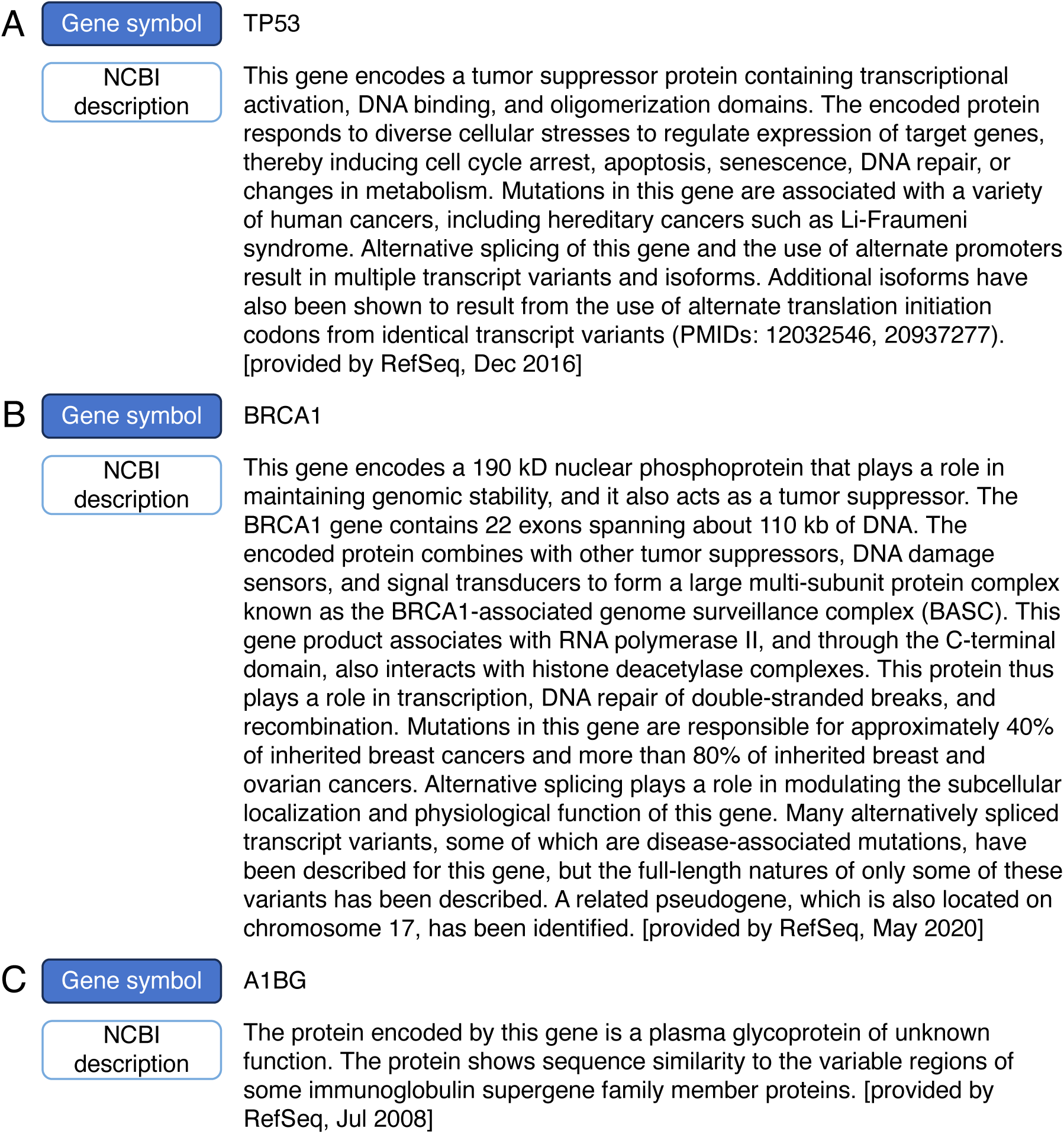
NCBI descriptions for three human genes: **(A)** *TP53* and **(B)** *BRCA1*, which are extensively studied yet described only briefly, and **(C)** *A1BG*, an example of an even shorter description typical for less-characterized genes.

This lack of detail is not unique to *TP53* and *BRCA1* but is, in fact, a common characteristic of gene descriptions in NCBI. Typically, the NCBI description of a gene is limited to one paragraph, containing no more than a few hundred words. For less-characterized genes, these descriptions are even more concise. As an illustration, Figure 1 (C) shows the description of the human gene *A1BG*, which appears first in the alphabetical list of human protein-coding genes [29]. The description of *A1BG* is very brief, consisting of only two sentences.

Given the brevity of these descriptions, it is not immediately evident or intuitive that LLM-generated gene embeddings derived from NCBI descriptions can capture comprehensive or even adequate information about genes, particularly their regulatory relationships and pathway associations. This inherent limitation raises critical questions about the extent to which such embeddings can encapsulate biologically meaningful information, warranting a rigorous evaluation.

### Pathway Analysis on OpenAI Text-Embeddings of Human Genes

We obtained the human gene embeddings generated by the GenePT study from its Zenodo repository. These embeddings were produced using OpenAI’s text-embedding-3-small model [30]. In generating each embedding, the input to the model was the official gene symbol concatenated with its NCBI description (examples shown in Figure 1), and the output was a 1,536-dimensional embedding vector, as shown in the top row of Table 1.

**Table 1:** Information on the OpenAI embedding model and the eleven small language models (SLMs) used in this study. Documentation for OpenAI embeddings is available at [30]. Information on the evaluated small language models, including training objectives and benchmark performance, is compiled from the MTEB benchmark and associated Hugging Face model documentation.

| Models | Model parameters<br>(million) | General-purpose | MTEB Average<br>Performance | Embedding<br>dimensions |
| --- | --- | --- | --- | --- |
| OpenAI - text-embedding-3-small | – * | Yes | 62.3 | 1536 |
| GIST-small-Embedding-v0 | 33 | Yes | 76.1 | 384 |
| NoInstruct-small-Embedding-v0 | 33 | Yes | 76.0 | 384 |
| stella-base-en-v2 | 55 | Yes | 75.3 | 768 |
| e5-small-v2 | 33 | Yes | 72.9 | 384 |
| GIST-all-MiniLM-L6-v2 | 23 | Yes | 72.7 | 384 |
| gte-small | 33 | Yes | 72.3 | 384 |
| bge-small-en-v1.5 | 33 | Yes | 72.2 | 384 |
| MedEmbed-small-v0.1 | 33 | Yes (partially **) | 71.8 | 384 |
| gte-tiny | 23 | Yes | 70.4 | 384 |
| e5-small | 33 | Yes | 68.1 | 384 |
| biobert-base-cased-v1.1 | 110 | No | – *** | 768 |

\* OpenAI does not disclose the parameter count or architectural details of this model.
\*\* MedEmbed-small-v0.1 is built on BGE-small and fine-tuned on synthetic medical QA data from PubMed Central, yielding improved biomedical performance while remaining broadly general-purpose and competitive on MTEB.
\*\*\* As BioBERT-base-cased-v1.1 is not intended as a general-purpose embedding model, it is not evaluated on MTEB, which encompasses a broad spectrum of task categories.

The primary objective of this study is to determine whether gene-regulatory pathway information is encoded within the high-dimensional gene embeddings. Our approach is straight-forward: for each of the 1,536 dimensions of the embeddings, we apply Gene Set Enrichment Analysis (GSEA) using pathways from the Human Molecular Signatures Database (MSigDB, [31, 32]). For each dimension, genes were ranked by their embedding coordinate values, and GSEA was applied to this ranked list. If a pathway is significantly enriched in a given dimension, we consider this as evidence that the dimension captures information about that pathway. We describe this as the pathway being “reflected” or “represented” by the dimension (not implying that the embedding dimension uniquely encodes this pathway). We further define “pathway coverage” as the proportion of pathways that are reflected by at least one embedding dimension:

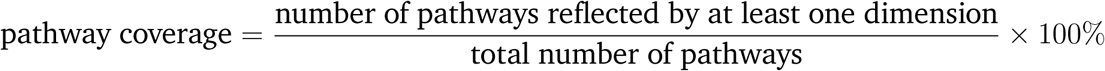

Pathway coverage serves as our primary evaluation metric. A higher pathway coverage suggests that the embeddings capture a broader range of pathway-level biological information. This per-dimension analysis allows us to identify which aspects of the embedding space align with known biological processes.

For GSEA, we will focus on two widely used collections of gene sets from the Human MSigDB: the 50 hallmark gene sets and the 3,917 canonical pathways in “C2” (curated gene sets). The hallmark gene sets [33] are coherently-expressed signatures derived by integrating many MSigDB gene sets to represent the most well-defined biological states or processes. In contrast, the 3,917 canonical pathways in C2 are curated from various sources, including online pathway databases and the biomedical literature. These pathways encompass six main categories: 292 BioCarta, 658 KEGG Medicus, 196 PID, 1,736 Reactome, 830 WikiPathways, and 186 KEGG Legacy gene sets. An additional 19 pathways classified as “miscellaneous” are excluded from our analysis.

Further, we retain only the protein-coding genes [29] within each gene set and remove any gene sets containing fewer than 10 such genes, as sets with fewer than 10 genes are unlikely to show significant enrichment due to limited statistical power. After this filtering step, all 50 Hallmark gene sets and 3,077 C2 canonical pathways (220 BioCarta, 398 KEGG Medicus, 196 PID, 1,367 Reactome, 711 WikiPathways, and 185 KEGG Legacy) are retained for further analysis.

GSEA is performed using the R package Fast Gene Set Enrichment Analysis (FGSEA) [34], which allows for false discovery rate (FDR) estimation and control at a user-specified threshold. The analysis is conducted independently for each of the 1,536 embedding dimensions, and pathway coverage is then computed by aggregating the statistically significant pathways across all dimensions. To control the overall false positive rate, we adopt a stringent multiple-testing correction strategy inspired by the Bonferroni method: for each embedding dimension, we set the FDR cutoff to *p/d*, where *p* = 0.05 and *d* is the dimension of the embedding. This conservative strategy is intended to minimize false discoveries and avoid overestimating the extent of pathway-level information captured by the embeddings. We choose this highly conservative threshold to avoid inflating pathway coverage due to false positives.

## Results

### Results on the Hallmark and C2 Gene Sets

Of the 50 Hallmark gene sets, 49 are represented by at least one embedding dimension, giving a pathway coverage of 98%. Furthermore, as shown in Figure 2(A), all but one of these significant gene sets are represented by more than one embedding dimension, with an average of 84.36 embedding dimensions per pathway. This redundancy indicates that pathway-level information is distributed across multiple dimensions rather than isolated within a single axis of the embedding space. The only Hallmark gene set not covered by any dimension is HALLMARK_NOTCH_SIGNALING (the prefix “HALLMARK_” has been omitted from all gene set names in Figure 2(A) to save space), which contains only 32 protein-coding genes—the smallest number among all Hallmark gene sets—whereas all other Hallmark sets contain at least 36.

**Figure 2:**
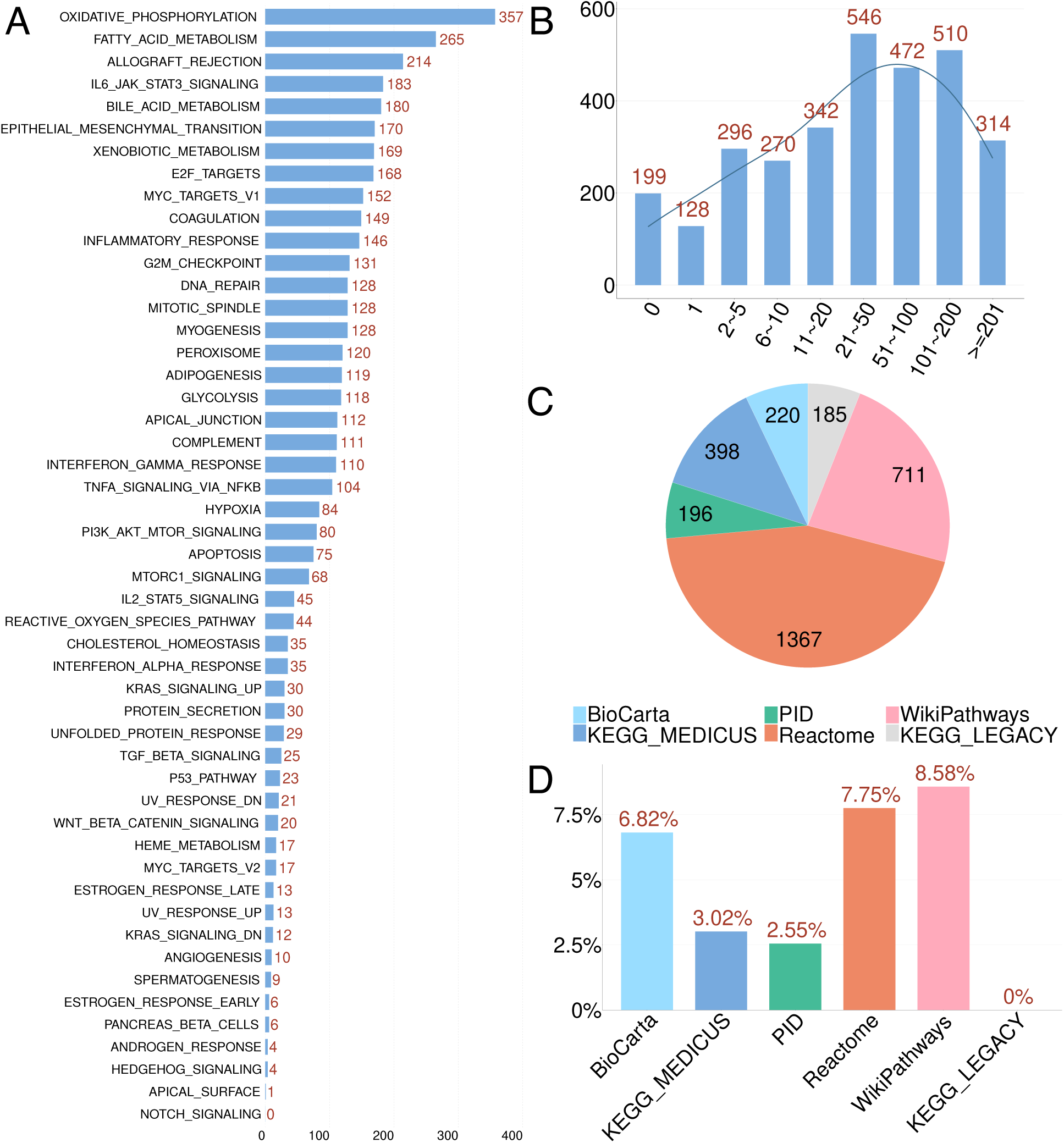
Pathway coverage of OpenAI gene embeddings. **(A)** Number of embedding dimensions in which each Hallmark pathway is detected as significant. **(B)** Distribution of the number of embedding dimensions representing each C2 pathway. **(C)** Total number of C2 pathways in each of the six MSigDB categories. **(D)** Percentage of pathways within each category that are not detected as significant.

We observe similar results for the C2 canonical pathways. Of the 3,077 C2 pathways, 2,878 are represented by at least one embedding dimension, yielding a pathway coverage of 93.53%. Most (95.55%) of them are represented by more than one dimension, with an average of 73.52 dimensions per pathway. Figure 2(B) shows the distribution of the number of embedding dimensions representing each C2 pathway.

C2 pathways can be further divided into six categories, with their respective sizes shown in Figure 2(C). Figure 2(D) displays, for each category, the percentage of pathways that are not detected as significant. Three categories—BioCarta, Reactome, and WikiPathways—show similar non-detection rates (6–9%). By contrast, KEGG_MEDICUS exhibits a lower rate of 3.02%, PID shows 2.55%, and KEGG_LEGACY shows no non-detected pathways, with all 185 pathways detected as significant.

The variability across categories is interpretable in light of known differences in pathway curation and biological coherence. KEGG_LEGACY and PID consist largely of well-established, mechanistically coherent pathways that are extensively represented in the biomedical literature [35, 36]; such pathways are more likely to exhibit consistent structure in the embedding space and thus be detected by our analysis. In contrast, Reactome and WikiPathways include many hierarchically organized or community-curated pathways whose granularity and gene-set coherence vary widely [37, 38], which can dilute enrichment signals and lead to higher non-detection rates. BioCarta, an older and partially deprecated collection, also contains pathway definitions that vary in specificity and are less consistently used in contemporary analyses [31]. These differences in curation style, pathway stability, and literature representation likely contribute to the observed variation in pathway detection across categories.

To summarize, the very high pathway coverage clearly indicates that the OpenAI gene embeddings contain rich biological information—not only for the relatively small and well-curated Hallmark gene sets, but also for the much larger and more heterogeneous C2 canonical pathways collected from diverse sources.

### Quality Check: Results on a Completely Randomized Embedding Matrix

As mentioned in the Methods section, we employed a stringent strategy to control false discoveries by using a very small FDR cutoff for each embedding dimension. Here, we conducted an empirical assessment to evaluate the actual rate of false discoveries. We randomly shuffled all elements of the *n*×*p* embedding matrix—where *n* is the number of genes and *p* is the embedding dimension—and repeated GSEA using the same correction strategy. On this shuffled matrix, the pathway coverage should ideally be close to zero if the multiple-testing correction is working properly.

Indeed, the observed pathway coverage on the shuffled matrix was 0.00% for both the Hallmark gene sets and C2 pathways. In other words, there is no pathway found to be significant, regardless of pathway collection. This result suggests that the false discovery rate is indeed controlled at an extremely low level.

### Assessing the Impact of Using Less Informative Text to Generate Embeddings

The near-perfect pathway coverage suggests that the OpenAI embeddings derived from NCBI gene descriptions encode a substantial amount of pathway-level information. This result is remarkable, given that NCBI gene descriptions are typically concise and contain limited explicit information about gene–gene regulatory relationships.

Why might such extensive pathway recovery occur? One possibility is that OpenAI’s text embedding model possesses exceptionally strong capabilities—potentially beyond what humans can extract from such brief descriptions. Such understanding could arise from the model’s ability to compare and contrast descriptions across many genes.

Another possibility is that the embeddings are not generated solely from the input text (i.e., the NCBI gene description), but rather represent a synthesis of the input and the model’s prior knowledge acquired during pre-training. As a transformer-based model trained on massive, diverse language corpora—including extensive biomedical literature—OpenAI’s embedding model may have been exposed to much richer and more explicit biological content than what is available in the NCBI descriptions. This background knowledge could be implicitly incorporated into the embeddings, even when the input text is limited.

While the first hypothesis is difficult to test directly and is left for future investigation, we designed an experiment to probe the second hypothesis. Specifically, we generated new embeddings for each gene using only the official gene symbol as input to the OpenAI embedding tool. We then performed the same GSEA analysis on these “symbol-only” embeddings and compared the results to those obtained using the full NCBI descriptions (hereafter referred to as “full-description” embeddings). To complete the comparison, we also evaluated an intermediate case: embeddings generated using the gene symbol concatenated with only the first half of the NCBI description (defined as the first 50% of words in the description), omitting the second half. We refer to these as “half-description” embeddings. In Figure 3(A), we illustrate the workflow used to generate the full-description, half-description, and symbol-only embeddings. Figure 3(B) provides a concrete example using the gene *TP53*, showing the exact text input used to produce each of the three embedding types.

**Figure 3:**
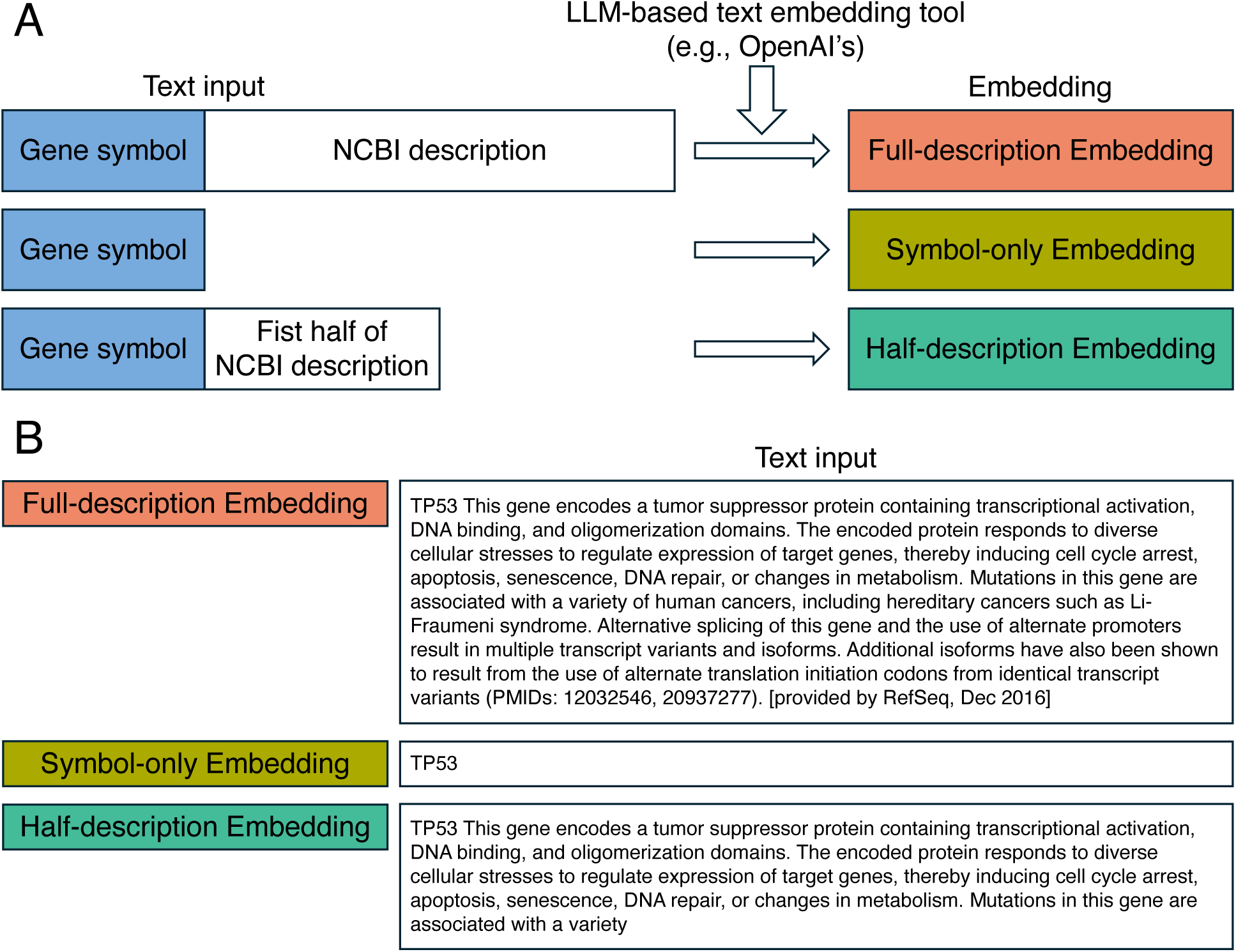
Generation of gene embeddings under different input settings. **(A)** Schematic illustrating the inputs used to generate the three embedding types: full-description (gene symbol + full NCBI description), symbol-only (gene symbol alone), and half-description (gene symbol + first half of the NCBI description). Each input is processed by an LLM-based text-embedding tool (e.g., OpenAI’s). **(B)** Example text inputs for the three embedding types, shown for the gene *TP53*.

The results on Hallmark pathways and C2 pathways are shown in Figure 4 (A) and (B), respectively. The symbol-only embeddings yielded pathway coverage of 64% (i.e., 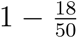) for the Hallmark gene sets and 72.15% (i.e., 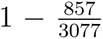) for the C2 pathways. While this coverage is notably lower than that of the full-description embeddings (98% and 93.53%, respectively), the half-description embeddings recovered most of this difference, achieving 96% (i.e., 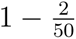) and 92.10% (i.e., 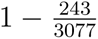). On the one hand, this confirms that the text descriptions contribute substantially to the captured pathway information. On the other hand, the fact that symbol-only embeddings alone achieve relatively high coverage suggests that a non-trivial portion of the pathway knowledge embedded in the OpenAI embeddings likely originates from the model’s prior biological knowledge acquired during pre-training.

**Figure 4:**
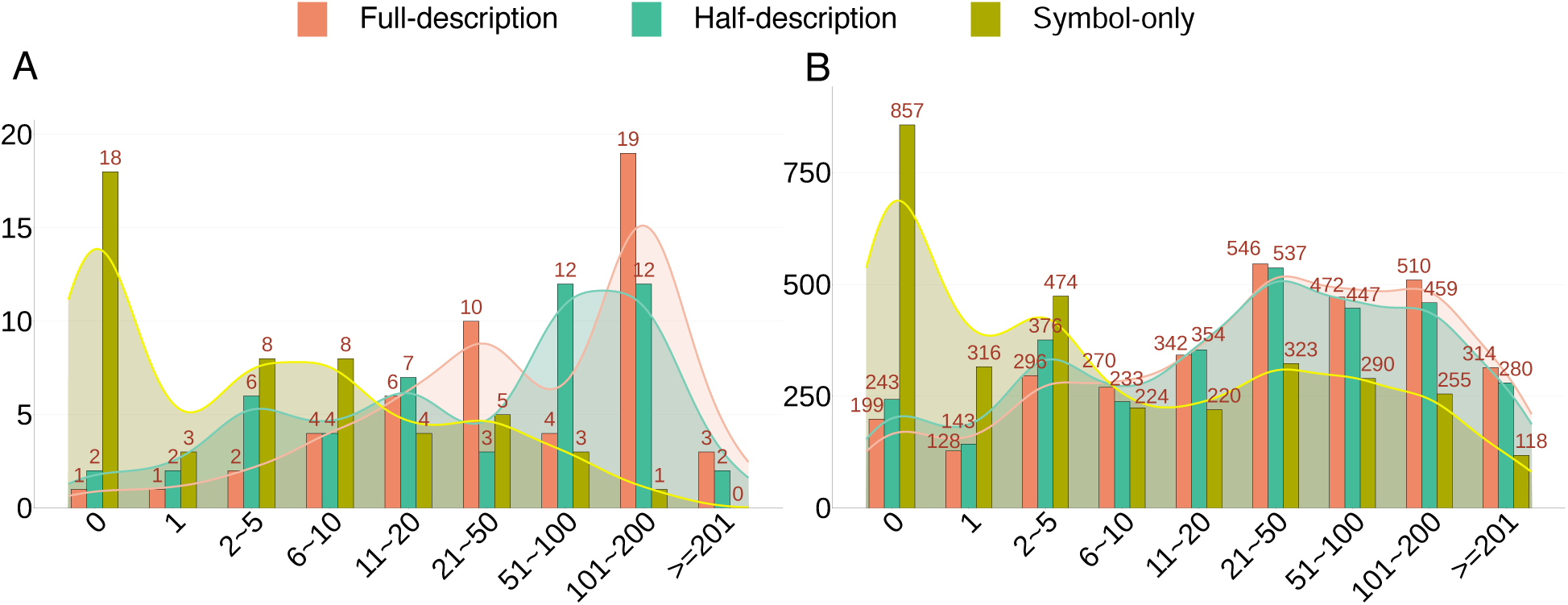
Distribution of pathway detection counts across embedding types. **(A)** Hallmark pathways. **(B)** C2 pathways. For each pathway, we record the number of embedding dimensions in which it is detected as significant, and group pathways into bins according to this count. Bar heights indicate the number of pathways in each bin. Colors correspond to embeddings generated from different input texts: full-description, half-description, and symbol-only.

### Assessing the Impact of Using Smaller LLMs to Generate Embeddings

To further investigate why OpenAI embeddings derived from NCBI gene descriptions encode such rich pathway-level information—despite the brevity of the input—we examined other LLM-based text embedding tools. Comparing these models may help shed light on the source of the information captured in the embeddings.

A recent study by Gan et al. [28] explored the use of embeddings generated by small language models (SLMs) for predictive tasks in gene-expression analysis. Specifically, the study evaluated ten open-source SLMs—listed from the second to the eleventh row in Table 1 (i.e., all rows except the first and the last)—each containing only 23 to 55 million parameters. For context, OpenAI’s GPT-3 has 175 billion parameters—over a thousand times larger—and GPT-4 reportedly contains 1.76 trillion parameters.

All ten models in the study are general-purpose, with one partial exception: MedEmbed-small-v0.1. This model is based on the general-purpose BGE-small model but was fine-tuned using contrastive learning on synthetic medical question–answer triplets derived from PubMed Central notes. While this fine-tuning improves its performance in clinical and biomedical tasks, MedEmbed-small-v0.1 remains usable as a general-purpose model and demonstrates strong performance on the Massive Text Embedding Benchmark (MTEB) [39].

Gan et al. [28] found that embeddings generated by these SLMs from NCBI gene descriptions performed comparably to OpenAI embeddings in various predictive tasks, including the prediction of long-range transcription factors, dosage-sensitive genes, and bivalent chromatin states. This naturally raises a question: Do these smaller models also capture similar amounts of gene-pathway information?

To explore this, we additionally included a domain-specific SLM, BioBERT-base-cased-v1.1 (last row in Table 1), in our analysis. Unlike the ten SLMs considered by Gan et al. [28], BioBERT contains 110 million parameters and is pre-trained on domain-specific biomedical corpora, including 4.5 billion words from PubMed abstracts and 13.5 billion words from PMC full-text articles [40]. It is considered a domain-specific model rather than a general-purpose one.

Figure 5 shows the pathway coverage of embeddings generated by these 11 SLMs, as well as by OpenAI’s model, when applied to the full NCBI gene descriptions. It also includes the results for the symbol-only and half-description embeddings. The rows of the figure correspond to different models, ordered by their average pathway coverage across all six conditions (two pathway sets × three input types). The name of the only domain-specific model, biobert-base-cased-v1.1, is shown in red, whereas the partially domain-adapted model, MedEmbed-small-v0.1, is shown in green. Several interesting findings emerge from this figure; we summarize them here and discuss them in greater detail in the Discussion section.

**Figure 5:**
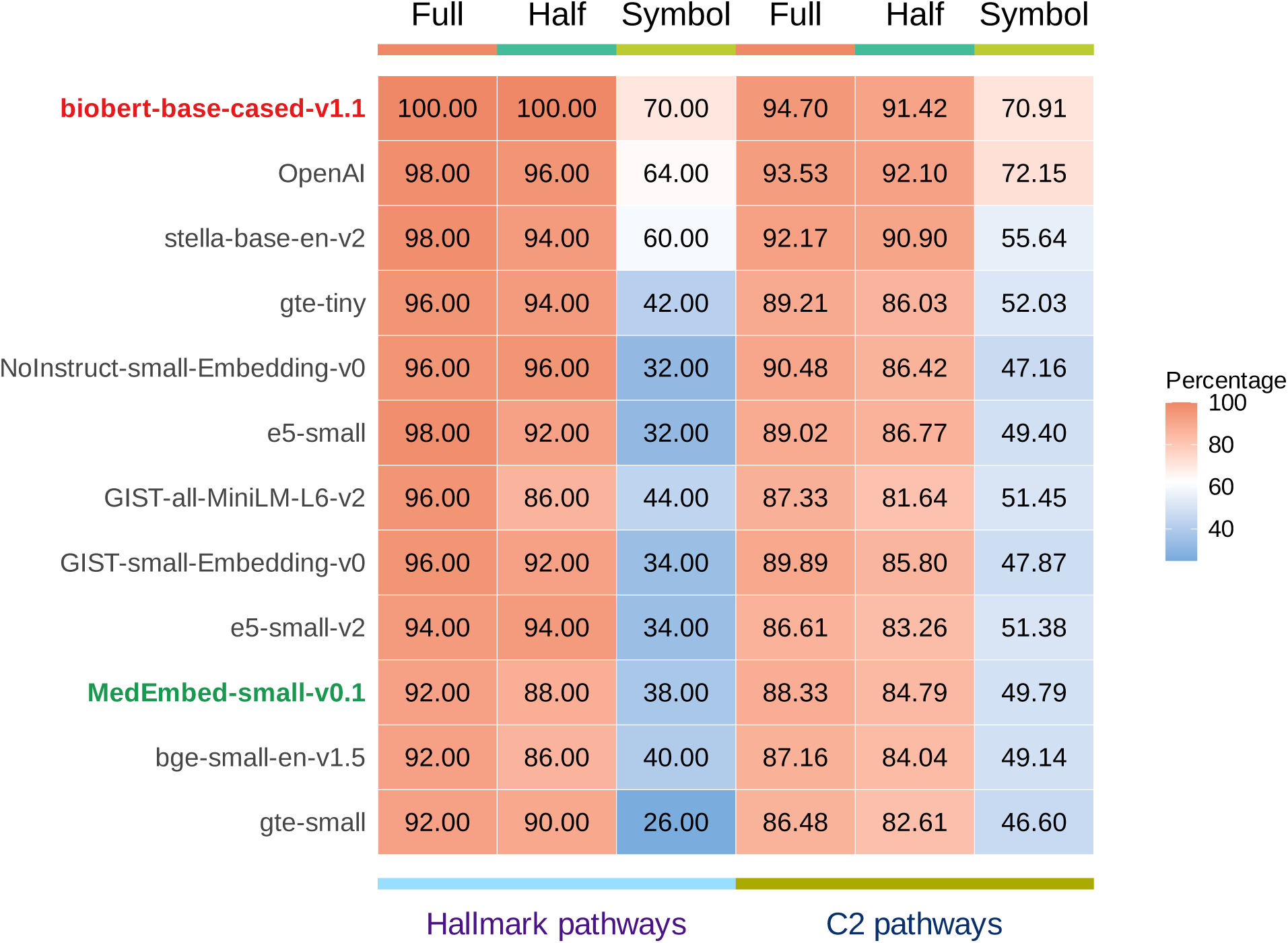
Pathway coverage of embeddings generated by different language models. Shown are OpenAI’s embedding model, nine general-purpose small language models (SLMs), one partially domain-adapted SLM (model name shown in green), and one domain-specific biomedical SLM (model name shown in red). For each model, three types of embeddings are generated: full-description (“Full”), half-description (“Half”), and symbol-only (“Symbol”). The left three columns report pathway coverage for the Hallmark pathways, and the right three columns report coverage for the C2 pathways. Higher percentages indicate that a greater proportion of pathways are detected as significant by at least one embedding dimension.

#### Four observations stand out

First, there is considerable variation in the pathway coverage of symbol-only embeddings across models. For Hallmark pathways, coverage ranges from 26% to 70%; for C2 pathways, from 46.60% to 72.15%. Most SLMs exhibit substantially lower coverage than OpenAI’s embeddings. However, BioBERT-base-cased-v1.1 achieves the highest coverage on the Hallmark pathways—even exceeding OpenAI’s (70.00% vs. 64.00%)—and ranks second on C2 pathways, with coverage nearly matching that of OpenAI (70.91% vs. 72.15%).

Second, in contrast, MedEmbed-small-v0.1, which is fine-tuned on clinical and medical texts, performs relatively poorly on both Hallmark and C2 pathways in the symbol-only setting. This suggests that fine-tuning on clinical data may not confer an advantage for capturing gene-regulatory information from gene symbols alone.

Third, it is notable that the small general-purpose model stella-base-en-v2 performs surprisingly well. Its symbol-only embeddings achieve relatively high pathway coverage, especially for the Hallmark pathways, outperforming all other small models except BioBERT-base-cased-v1.1. For the C2 pathways, both their symbol-only and full-description embeddings outperform those of all models except BioBERT-base-cased-v1.1 and OpenAI.

Fourth, when full NCBI gene descriptions are used, the pathway coverage of all models increases substantially and approaches that of OpenAI’s embeddings. Specifically, the pathway coverage from full-description embeddings exceeds 86% for all models on both Hallmark and C2 pathways. Even with only half of the NCBI description (half-description embeddings), coverage remains above 81% for all models and all kinds of pathways. These findings suggest that the performance gap between small language models and OpenAI’s model narrows quickly with the inclusion of modest textual context from the gene description. This provides further support for the hypothesis that the models extract implicit and relational cues from even highly compressed text.

## Conclusions and Discussion

OpenAI’s gene embeddings, generated from NCBI gene descriptions, have demonstrated exceptional predictive performance in various biological applications. However, it remains unclear how much biologically meaningful information—particularly pathway-level knowledge—is actually encoded in these embeddings. Moreover, if such rich information is indeed present, it raises the intriguing question of how this is possible, given the brevity and limited detail of the NCBI gene descriptions. In this study, we provide an initial exploration of these questions.

Using a GSEA-based framework, we found that OpenAI’s embeddings achieve pathway coverage exceeding 93% for both Hallmark and C2 pathways. Considering the broad and heterogeneous nature of the C2 pathways, this level of coverage is striking. It indicates that a vast majority of pathway-level information is captured by these embeddings, suggesting that they are well suited for downstream applications involving gene–gene interactions and pathway inference. Furthermore, we observed that even the symbol-only embed-dings—generated using just the gene symbol—capture over 64% of the Hallmark and C2 pathways. This result points to the substantial prior biological knowledge retained in the OpenAI embedding model, likely acquired during pretraining.

To better understand why OpenAI’s embeddings capture such a vast amount of pathway-level information—despite being derived from relatively short gene descriptions—we examined embeddings generated by small language models (SLMs), which differ markedly from OpenAI’s model in both scale and pretraining data. While the Results section outlined four key empirical findings from this comparison, here we reflect on what these findings reveal about the underlying factors contributing to pathway-level knowledge in different embedding models.

First, the strong performance of the symbol-only embeddings from BioBERT-base-cased-v1.1 is expected. This model is the only one in our study that was pretrained exclusively on a large corpus of biomedical text. As a result, it has learned domain-specific representations of genes and their regulatory functions. Thus, even when only given a gene symbol, the model can retrieve relevant information from its internal parameters and encode it into embeddings. While BioBERT has far fewer parameters than OpenAI’s model, its specialized training appears to confer a clear advantage for this task.

Second, the relatively poor performance of MedEmbed-small-v0.1 is also understandable. Although it was fine-tuned on medical text, its training data—clinical note–derived synthetic triplets—are primarily patient-facing or oriented toward clinical trials. Consequently, it is less well-suited for capturing molecular or pathway-level gene information.

Third, the surprisingly strong performance of stella-base-en-v2 may be attributed to several factors. It is the largest among the general-purpose SLMs evaluated (55 million parameters) and was trained on a large and diverse corpus (∼200 GB) that includes long-form, knowledge-rich sources such as Wikipedia and web articles, many of which likely contain general biological content [41]. Additionally, it was trained using contrastive learning with hard negatives [42] and LLM-generated question–paragraph pairs [41], some of which may implicitly encode biomedical facts. The model’s architecture and training design—including document-level training and absence of instruction prefixes—may further enhance its ability to generalize factual knowledge across domains [43]. Taken together, these characteristics distinguish stella-base-en-v2 from other general-purpose SLMs evaluated in this study, many of which are constrained by design choices that prioritize retrieval-oriented objectives over factual recall or domain generalization (Table 2).

**Table 2:** Likely factors contributing to the underperformance of the eight evaluated general-purpose SLMs (excluding stella-base-en-v2) on biology-related embedding tasks.

| Model | Likely Limitation | Reference |
| --- | --- | --- |
| bge-small-en-v1.5 | Trained primarily for general-purpose semantic retrieval using contrastive loss; not optimized for factual recall or scientific domain generalization | [48] |
| e5-small / e5-small-v2 | Focused on instruction-tuned embedding for search and QA; not exposed to biomedical corpora | [49] |
| GIST-all-MiniLM-L6-v2 | Designed for dense passage retrieval and knowledge-intensive QA, not factual recall or domain coverage | [50] |
| GIST-small-Embedding-v0 | Optimized for semantic similarity and long document structure, not factual or domain-specific content | [51] |
| gte-small / gte-tiny | Extremely compact; trained for retrieval-style tasks using MTEB and general-purpose corpora | [52] |
| NoInstruct-small-Embedding-v0 | Trained without instruction formatting; optimized for instruction-agnostic retrieval rather than domain knowledge | [53] |

Fourth, the observation that all SLMs—despite their much smaller size—approach OpenAI’s performance when given the full NCBI gene description (or even just half of it) suggests that modern LLMs are highly effective at extracting both explicit and implicit information from short texts. Their ability to discern nuanced relationships and semantic signals from limited context plays a critical role in embedding quality.

Taken together, these findings provide insight into a central question: Do LLM-derived gene embeddings derive their apparent knowledge from the textual input (i.e., NCBI descriptions) or from the model’s internalized prior knowledge? Our results suggest that both factors contribute, although the relative contribution likely varies across models depending on domain specialization and corpus composition. Different LLMs encode different levels of pathway knowledge, based on their pretraining corpora. At the same time, all models tested in this study demonstrate strong capabilities for inferring and extracting biological information even from brief textual inputs.

Despite the simplicity of our GSEA-based approach—treating each embedding dimension as a potential carrier of pathway information—it has a noteworthy limitation: pathway coverage is not invariant under rotations of the embedding space. Formally, let *A* be an *n* × *p* embedding matrix, where *n* is the number of genes and *p* is the embedding dimension. Let *R* be any *p* × *p* orthogonal matrix such that *R*^⊤^*R* = *RR*^⊤^ = *I*. Then the transformed matrix *Ã* = *AR* preserves all pairwise relationships and geometric structure in the embedding space. However, since GSEA is conducted on individual columns of *A* (i.e., specific dimensions), the pathway coverage depends on the choice of *R*, which has infinitely many possibilities. That is, *A* and *Ã* contain the same total information, yet they may yield different coverage scores. Thus, pathway information encoded by an LLM-generated embedding matrix should, in principle, be measured by the maximum pathway coverage achievable over all possible rotations *R*.

With this in mind, we devoted substantial effort to identifying an optimal rotation *R*. We adopted methods from the Crawford-Ferguson (CF) rotation family [44], commonly used in factor analysis to improve interpretability. Inspired by the “word intrusion” task in natural language processing, we aimed to rotate the embedding matrix to a form that yields a more interpretable and sparse structure. The rotation was obtained by minimizing the following cost function:

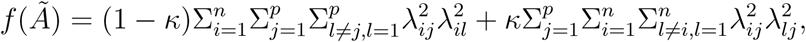

where *λ_ij_* is the (*i, j*)-th entry of the rotated embedding matrix *Ã*, and *κ* ∈ [0, 1] is a hyper-parameter balancing row sparsity (gene-level specificity) and column sparsity (dimension-level specificity). Optimization was performed using the gradient projection method [45, 46], and a GPU-accelerated implementation from [47].

We experimented with four commonly used *κ* values—corresponding to the Quartimax, Varimax, Parsimax, and FacParsim criteria—and ultimately selected Parsimax based on the “distance ratio” metric. We also considered a relaxed version of the problem where *R* is not required to be strictly orthogonal, but this variant yielded poorer distance ratios and was not adopted.

Despite these efforts, we found that the rotated embeddings *Ã* = *AR* did not lead to improved pathway coverage. Specifically, the rotated matrix yielded 94.00% and 92.79% coverage for Hallmark and C2 pathways, respectively—slightly lower than the original matrix (*A*), which achieved 98.00% and 93.53%. This suggests that, although rotation may improve interpretability in some contexts, it offers limited benefit in maximizing pathway representation in our setting. Ultimately, the rotation procedure is an unsupervised optimization task, and the objective function does not incorporate the biological goal of pathway enrichment. As such, the rotated embedding space may be suboptimal for our specific purpose.

Nonetheless, our attempt at rotation is carefully documented in the Supplementary Materials, including methodology, implementation details, and results. We believe this documentation will be valuable for researchers seeking to improve the interpretability of gene embeddings or apply similar techniques in other biological contexts.

Importantly, the lack of improvement through rotation does not diminish the conclusions of this study. From a theoretical standpoint, our reported pathway coverage—based on the unrotated embedding matrix—is a lower bound on the true pathway coverage achievable under optimal rotation. Thus, the true extent of biologically meaningful information encoded in these embeddings may be even higher than what we report here.

Collectively, our results demonstrate that LLM-derived gene embeddings encode unexpectedly extensive pathway-level information, highlighting their potential as a lightweight yet powerful resource for downstream biological analysis and motivating deeper methodological exploration.

## Data availability

All datasets used in the study have been published and cited in the main body.

